# Balancing postural control and motor inhibition during gait initiation

**DOI:** 10.1101/2024.05.16.594550

**Authors:** Lorenzo Fiori, Surabhi Ramawat, Isabel Beatrice Marc, Valentina Giuffrida, Alberto Ranavolo, Francesco Draicchio, Pierpaolo Pani, Stefano Ferraina, Emiliano Brunamonti

**Affiliations:** Department of Occupational and Environmental Medicine, Epidemiology and Hygiene, INAIL, Rome, Italy; Behavioral Neuroscience PhD Program, Sapienza University, 00185, Rome, Italy; Department of Physiology and Pharmacology, Sapienza University, 00185, Rome, Italy

**Keywords:** *Gait initiation*, *Postural control*, *Motor inhibition*, *Stop Signal Task*

## Abstract

This study examines the relationship between stopping a planned gait initiation due to sudden environmental changes and maintaining body stability. Using a gait initiation version of the Stop Signal Task (SST), we studied changes in anticipatory postural adjustments (APA) during gait initiation and suppression. We found that trial-level variables, such as the time to start or stop stepping, interacted with biomechanical factors like the center of mass displacement relative to the base of support, affecting performance. A critical biomechanical threshold was identified, beyond which stopping movement was unlikely. These findings highlight the strong link between limb action control and body equilibrium, offering a framework within a motor control paradigm. By integrating biomechanical elements, the model effectively simulates real-life scenarios, identifying key variables for studying neural correlations between action and postural control, and aiding in the development of injury prevention and rehabilitation tools for individuals with movement and posture impairments.

## Introduction

A primary purpose of posture control is to preserve body stability during voluntary movements and environmental interactions ^1–5^. Gait, as full-body movement, has been described as a ‘controlled falling’ (Perry, 1992) and this is especially true during gait initiation when the motor system must cope with the challenge of safely translating the body from a stable stationary state to a new one ^6,7^. Before the first step, to ensure stability, the nervous system executes a set of anticipatory postural adjustments (APA) to avoid perturbations dangerous for stability.

This already taxing sequence can become even more challenging when abrupt demands emerge from the environment. For instance, when a pedestrian is moving after the green traffic light but a car speeds trough the crossway. In this situation, the APA anticipating the movement need to be interrupted and the motor system to deal with both cancelling the movement and activate a process that guarantee the maintenance of the body equilibrium ^8,9^. Thus, preventing falls during gait initiation is a complex process involving APA and requiring the evaluation of environmental and proprioceptive stimuli ^10–14^. While this dynamic has been widely studied, how APA are adjusted if the initial step cancellation is required it is still unknown.

Here we approached this problem by implementing a gait version of the Stop Signal Task (SST) focusing on the initial phases of control. The SST is a widely used paradigm that has provided concrete quantifications of several parameters of motor inhibition ^15,16^. These measures have been of relevance for evaluating the behavioral and physiological correlates of action control of simple movements ^16–18^. In a typical SST experimental setup, participants are instructed to start a movement in response to a Go signal, and to refrain from the movement if a Stop signal is presented during the reaction time ^16,19–24^. By referring to a race model, where a go and a stop process run one against the other toward a finish line, the outcome in trials requiring inhibition can be accounted as the process that crossed this line first ^15^. In this context, stopping a planned movement is easiest in trials with longer Go reaction time when the Stop signal occurs sufficiently before the movement onset or the reaction time to the Stop signal is fast. Different combinations of these variables could define the outcome of movement inhibition, with corresponding failures and successes.

If this model defines a general framework of motor inhibition, here we expect to observe these variables to determine when APA will be completed or cancelled. The recent observation that the reaction time to a stop signal computed for a finger keypress correlates with suppression of a balance recovery step ^25^ point towards this direction, but direct evidence is needed.

We tested this hypothesis by successfully measuring, at the single trial level, APA responses to both Go and Stop signals presented during gait initiation and highlighted that all the derived variables align well with the established framework of the race model. It provides valid evidence that this framework of motor control could be extended to naturalistic movements, as recently proposed ^16^.

The present findings hold significant applied value. For example, in the context of healthcare for vulnerable individuals, such as elderly or Parkinsonian patients, framing the observed gait initiation parameters within a theoretical context could aid in planning specific therapeutic interventions and monitoring their efficacy ^26^.

## Results

### APA changes capture the key parameters of gait initiation and inhibition at the single trials level

We asked 12 subjects to stand in a comfortable position in front of a monitor where visual stimuli were presented (Figure 1A). In most of the trials (Go trials, 70%), subjects were instructed to start walking using the right leg soon after the presentation of a Go signal. We considered the first step initiated when the heel released the contact from the floor. In a subset of trials (Stop trials, 30%), a Stop signal was presented at variable intervals (stop signal delay; SSD) from the Go signal. Successful Stop trials (Stop Correct) were characterized as the trials in which the subjects did not interrupt the contact with the floor and cancelled the planned step, while trials in which the step movement was initiated, after the presentation of the Stop signal, were classified as Stop Error trials. A staircase procedure, monitoring the behavioral outcome for each SSD value proposed, was used to ensure each subject had approximately 50% successful Stop trials (see Star Methods). We observed that in a subgroup of the Stop Error trials the foot position was rapidly restored after the initial heel release. We labelled these trials as Stop partial Error trials (see Star Methods), as previously suggested in other contexts ^20,21,27^ .

**Figure 1.**
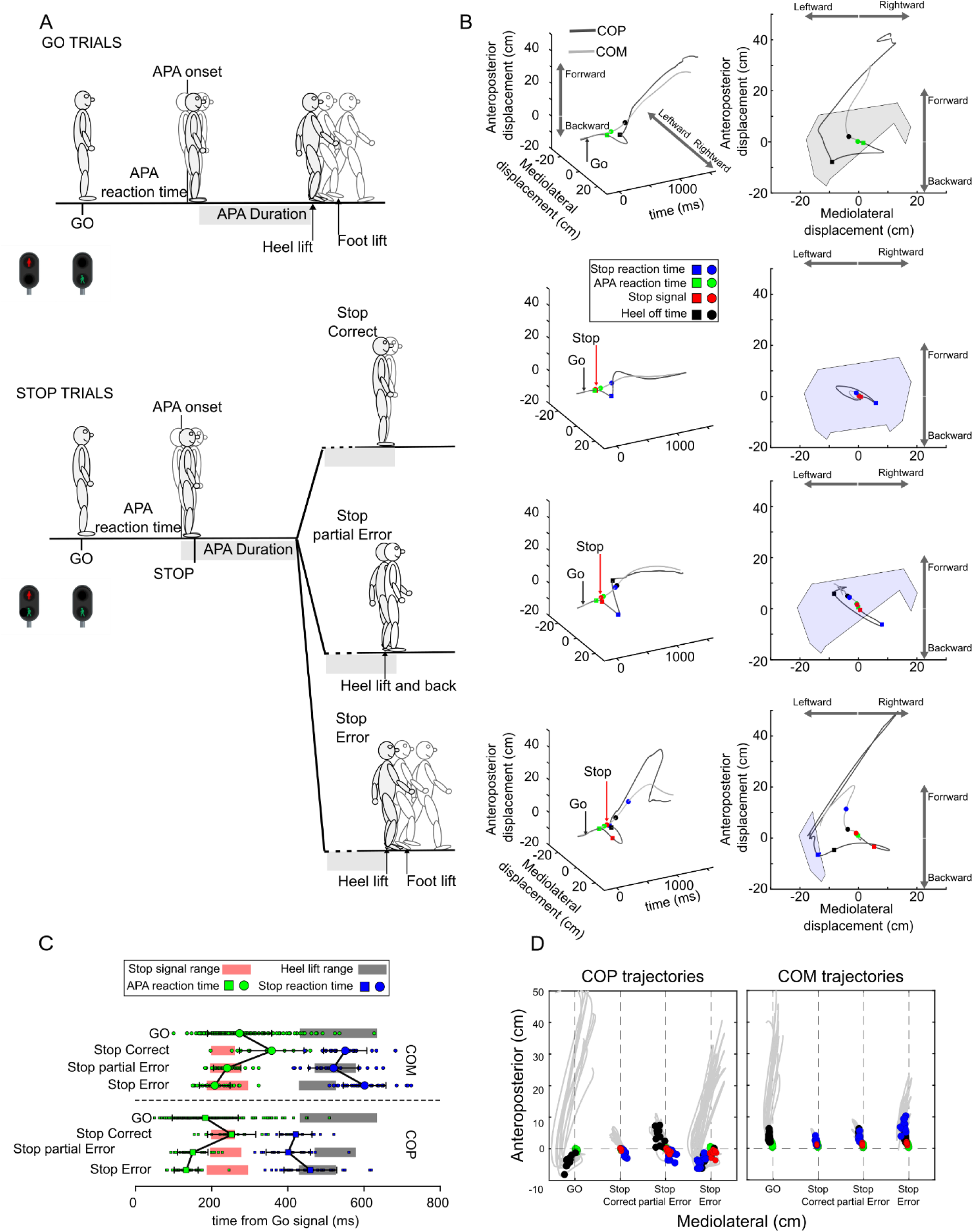
Illustration of the gait initiation stop signal task and trial by trial time evolution of Center of pressure (COP) and center of mass (COM) signals. A) A schematic of the behavioral outcome in the different task trials B) time course of the Mediolateral and Anteroposterior shifting of a single trial of the four possible outcomes in the task of a participant. Green squares and circles mark for each trial the time of APA onset obtained by the COP and COM respectively. In each stop trial, the time of Stop signal (red squares and arrows) and the corresponding reaction time (blue squares and circles) are indicated. The heel off time (dark square) in Go, Stop partial Error and Stop error trials are reported. Right panels also illustrate the extension of the BOS in Go trials at the time of heel lift (black area) and in Stop trials, at the time of the reaction rime to the Stop signal (light blue area). C) Displays the across trials distribution of the APA reaction time and the Reaction time to the Stop signal for the representative participant in the different types of trials of a single participants. Average and s.e.m. of the different distribution is over-imposed to the distribution. Shaded grey areas mark the edges of the distribution of the heel lift time in Go, Stop partial Error and Error trials. Shaded pink areas marks the edge of distribution of the time of Stop signal onset in Stop trials. D) Illustration of the trajectory of COP and COM averaged across trials for the different behavioral outcomes in the 12 participants. The average of APA reaction time, time of Stop signal occurrence and Reaction time to Stop signal are projected on the 2D spatial displacement of the different trajectories.

Before the heel lift, a sequential activation of APA elements was observed, first involving the center of pressure (COP) and then the center of mass (COM). Figure 1B-C illustrates the COP and COM trajectories and their reciprocal relationship in each of the different behavioral outcomes.

Figure 1B (left column) illustrates, in a three-dimensional space, the temporal evolution of the anteroposterior and mediolateral displacement of the COP and COM in an example trial of the different behavioral conditions, from a participant performing the task. In Go trials (top panels), consistent with existing literature on step initiation and completion, we first observed a lateral shift of the COP towards the side of the stepping foot (the right foot in this study). This was followed by a lateral shift towards the supporting foot, and subsequently, forward progression, initiating first heel and then whole foot lifting. The COM followed these COP adjustments by initially shifting laterally and then forward. The rightmost panels of Figure 1B illustrates the relationship between these variables projected onto a mediolateral-anteroposterior-plane. The bottom panels of Figure 1B demonstrate that the presentation of the Stop signal elicited a modification of the temporal evolution of COP and COM displacement, which varied across different outcomes of Stop trials. By monitoring the velocity variation of these biomechanical events (see Star Methods and Figure S2), we captured the time at which the COP and COM responded to the Go signal for movement initiation, marking the corresponding APA reaction time across all trial types. Additionally, by examining the change in COM and COP velocity profiles, we estimated the time at which APA were rectified in response to the Stop Signal (Reaction time to the Stop Signal). These adjustments were further evaluated in function of extension of the Base of Support (BOS) at the time of the APA Reaction time to the Stop Signal in the different Stop trials. The different Stop trials illustrated in Figure 1B are characterized by different lengths of Reaction time to the Stop signal, appearing shorter in Stop Correct and Stop partial Error trials compared to Stop Error trials. These changes were accompanied by a lowering of the extension of BOS. In the latter, the Reaction time to the Stop signal obtained from the COP (blue squares) occurs in a time very close to the time of heel lift (black squares), while that obtained from the COM (blue circles) occurs even later, projecting the COM outside the edges of the BOS. Once obtained these estimates from each trial, we investigated whether their distributions could account for the outcomes. Figure 1C lower illustrates for a participant the distribution of both the APA reaction time across the different trials—Go, Stop Correct, Stop Partial Error, and Stop Error— and that of the Reaction time to the Stop signal obtained by both the COP and COM trajectories during Stop trials. The figure confirmed that the reaction to both the Go and Stop Signal was first detected by the adjustments of COP (green and blue squares) and then by those of COM (green and blue circles). The figure also illustrates that the relationship between the APA onset time obtained by either the COP or COM was comparable and evidenced longer in the Go and Stop Correct trials respect to Stop Error and partial Error trials (green circles and squares). Similar relationship in the Reaction time to the Stop signal was observed in COP and COM between the different types of Stop trials. For these trials we observed longer Reaction to the Stop signal in Stop Error trials than in the Stop Correct and partial Stop Error trials (blue circles and squares). Figure 1 D illustrates on a mediolateral-anteroposterior projection plane of the COP and COM across trials average trajectories of each participant in the different trial types, which evidences a consistent pattern of displacement of the trajectories between the participants in the different types of trials.

To summarize, the different behaviors detected during the gait version of the SST were defined by a different COP trajectory that in turn was reflected in the time evolution of COM displacement. Accordingly, we observed that variations of the different patterns of trajectories that captured the reaction times to the Go and Stop signals at a single trial level occurred with different latencies in the different types of trials.

### Evaluation of the detected measures in the framework of the horse race model

The complex nature of a step initiation, involving a prolonged period of APA, offers a privileged window for observing motor variables supporting the framework of the Stop Signal paradigm developed through the study of ballistic or semi-ballistic movements ^19,22,28–32^. In all these experimental contexts, a behavioral measure of the Reaction time to the Stop signal, commonly referred as Stop Signal Reaction Time (SSRT), is difficult to be obtained at a single trial level because inhibition refrains the overt movements. For the same reason, it is not possible to detect a reaction time to the Go signal in Stop Correct trials. Only recently, the trial-by-trial estimate of the SSRT in a few experimental studies has been obtained ^20,21,31,33,34^.

Here, we were able to measure these variables at the single-trial level by analyzing COP and COM trajectories (refer to Figure 1B and STAR METHODS). We also tested whether these variables met the theoretical assumptions of the horse race model.

To this end, as a first step, we tested if our data satisfied the assumption of independence between the Go and Stop processes, hypothesized to run independently one from another after being triggered by a Go and a Stop signal, respectively ^15,17^. According to this model, the trials with the longer reaction times will be the ones that will be more likely inhibited by the occurrence of a stop signal (Logan and Cowan, 1984). Here we tested this assumption by comparing the COP and COM reaction time to the Go signal between Go and Stop trials.

Figure 2A shows that the longer APA (COP and COM, separately) reaction time to the Go signal were observed in the Stop Correct trials and Go trials, while the shorter reaction time was observed in both type of Stop Error trials. Moreover, the APA reaction times of the Stop partial Error trials were longer than those of Stop Error trials, a condition in line with the possibility to rectify a Stop trial initially started as Stop Error. This trend of the reaction times to the Go signal was similarly detected by considering either the time evolution of the COP (Figure 2A: left panel) or the COM (Figure 2A: right panel). Within each set of measures, the differences between the Go reaction times were statistically significant. A one-way Anova revealed a significant difference between the distributions of the Go signal reaction time of COP (**F(1,3)**= 128.02, ***p***<0.001, ***ηp***^2^ = 0.92) and COM (**F(1,3)**= 84.29, ***p***< 0.001, ***ηp^2^*** = 0.89). Specifically, for COP, the mean APA Reaction times were as follows: 253.23 ± 67.97 ms (Go), 279.41 ± 53.56 ms (Stop Correct), 208.27 ± 66.32 ms (Stop partial Error), and 186.94 ± 62.60 ms (Stop Error). For COM, the mean APA Reaction times were: 343.42 ± 66.36 ms (Go), 387.74 ± 55.33 ms (Stop Correct), 310.23 ± 61.28 ms (Stop partial Error), and 278.78 ± 67.60 ms (Stop Error). Bonferroni post-hoc comparison detected for both COP and COM significant difference between each reaction time to the others (all ***ps***< 0.001).

**Figure 2.**
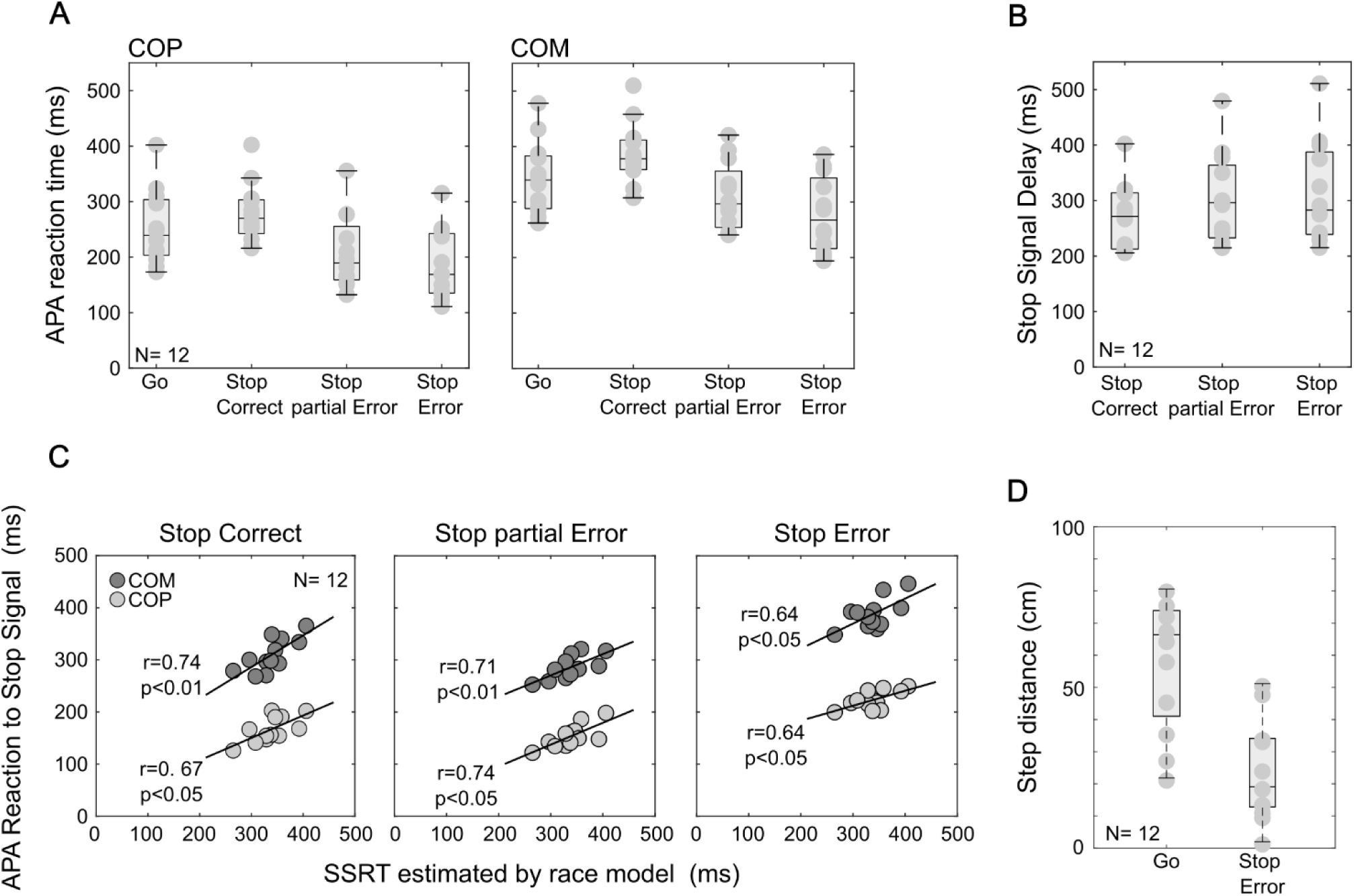
Stop signal task parameters estimated by the assumptions of the race model and the ones detected from the temporal evolution of the COP and COM. A. Boxplot comparing the APA reaction time to the Go signal between Go, Stop Correct, Stop partial error, and Stop error trials, detected in COP (left panel) and COM (right panel). B. Stop signal delay distribution in the different Stop trial categories for each considered parameter C. Correlation analysis between SSRT estimated by the race model and the APA reaction time to the Stop signal obtained by the kinetic and kinematic signals. D. Distance between initial and final step position in Go and Stop Error trials.

A further variable accounting for the Stop trials outcome in this framework is the link between the increased duration of the Stop Signal Delay (SSD) and the lower probability of cancelling prepared movements. Figure 2B illustrates that the length of SSD of Stop Correct trials was significantly different between the different Stop trials (one-way Anova: **F(1,2)**= 15.08, ***p***< 0.001, ***ηp^2^***= 0.57). Specifically, the mean SSDs were as follows: 269.86 ± 61.32 ms (Stop Correct), 302.83 ± 81.91 ms (Stop partial Error), and 312.21 ± 92.46 ms (Stop Error).

Bonferroni post-hoc comparisons detected that Stop correct SSD were significantly shorter than Stop partial error and Stop error trials SSD (all ***ps***< 0.05). However, no significant difference was detected between SSDs of the Stop Error and Stop partial Error trials (***ps***> 0.05).

Finally, we evaluated a further implication of the model that relates the SSRT to the outcome in Stop trials, a relationship that is rarely tested experimentally. In the context of the race model, the Reaction time to the Stop signal is expected to be longer in Stop Error trials than in Stop partial Error and Stop Correct trials. Considering the temporal resolution of our measures, here we were able to detect these variables in each different category of Stop trials (Figure 2C and Table 1). A one-way Anova confirmed that the observed differences were statistically significant (COP: **F(1,2)**= 118.02, ***p***< 0.001, ***ηp^2^*** = 0.92; COM: **F(1,2)**= 141.05; ***p***< 0.001, ***ηp^2^*** = 0.93; all Bonferroni post-hoc comparisons: ***ps***< 0.05).

**Table 1**
| Trial Type | COP | COM |
| --- | --- | --- |
| Stop Correct [ms] | 166.81(±24.83) | 310.02(±32.38) |
| Stop partial Error [ms] | 153.57(±21.88) | 286.37(±22.43) |
| Stop Error [ms] | 223.79(±18.32) | 390.00(±30.57) |

Computed APA Reaction time to the Stop signal in COP and COM. Mean (SD).

To verify that our measure of the APA Reaction time to the Stop signal complied with the theoretical horse race model, we correlated this variable, averaged across trials, with the SSRT estimated based on the model’s assumption and the related computational method (See Materials and Method). We observed a significant correlation between these two measures of inhibition (Figure 2C). For both COP and COM, we detected a significant correlation coefficient between the SSRT estimated by the race model’s assumptions and the one obtained by the APA displacement in the Stop Correct, Stop partial Error and Stop Error trials. This result is consistent with previous studies that identified a significant correlation between model-estimated SSRT and single-trial SSRT estimates obtained through comparable methods ^20,31,35^.

To further evaluate the dynamics of competition between the Go and Stop processes, we compared the stepping foot distance in Go trials (57.99 ± 20.09cm) and Stop Error trials (23.56 ± 15.41 cm). The analysis revealed a significantly smaller distance in Stop Error trials compared to Go trials (t-test: ***p*** < 0.001), indicating reduced movement between the initial and final foot positions during failed inhibition (Figure 2D). This subtends the occurrence the attempt of the stop process to interrupt the ongoing movement ^36^.

### Failure in canceling the step depends on the biomechanics of the body at the time of stop reaction

Our approach allowed us to detect, on the base of APA measures displacement, a Reaction time to the Stop signal, at single trial level, that significantly correlated with the behavioral estimate of the SSRT based on the race model assumptions as a reliable estimate of this parameter of inhibition. Since this reaction occurred in each of the different categories of Stop trials, here we asked when this ended up with a successful inhibition of the step initiation. To answer this question, we compared the area of the base of support (BOS; see Materials and Method) and the vertical projection of the COM on it (XCoM). Figure 3 shows that if at the time of the detected Reaction time to the Stop signal the XCoM fell within the area of BOS the step was successfully cancelled. This is the case of Stop Correct and Stop partial Error trials for which the inhibitory signal kept the body in the starting position (Figure 3 A-B). On the contrary, in Stop Error trials, the reduced area of BOS (Figure 3 A rightmost panel) did not allow the maintenance of the starting position. In these trials the loss of equilibrium was prevented by a forward step that increased the area of BOS (not shown). Across participants, we detected the antero-posterior and mediolateral displacement of COM (measured as Stability index; see Materials and Methods) in Stop Error trials was approximately 3 times, and significantly different, to that of Stop Correct and Stop partial Error trials (Figure 3 B).

**Figure 3.**
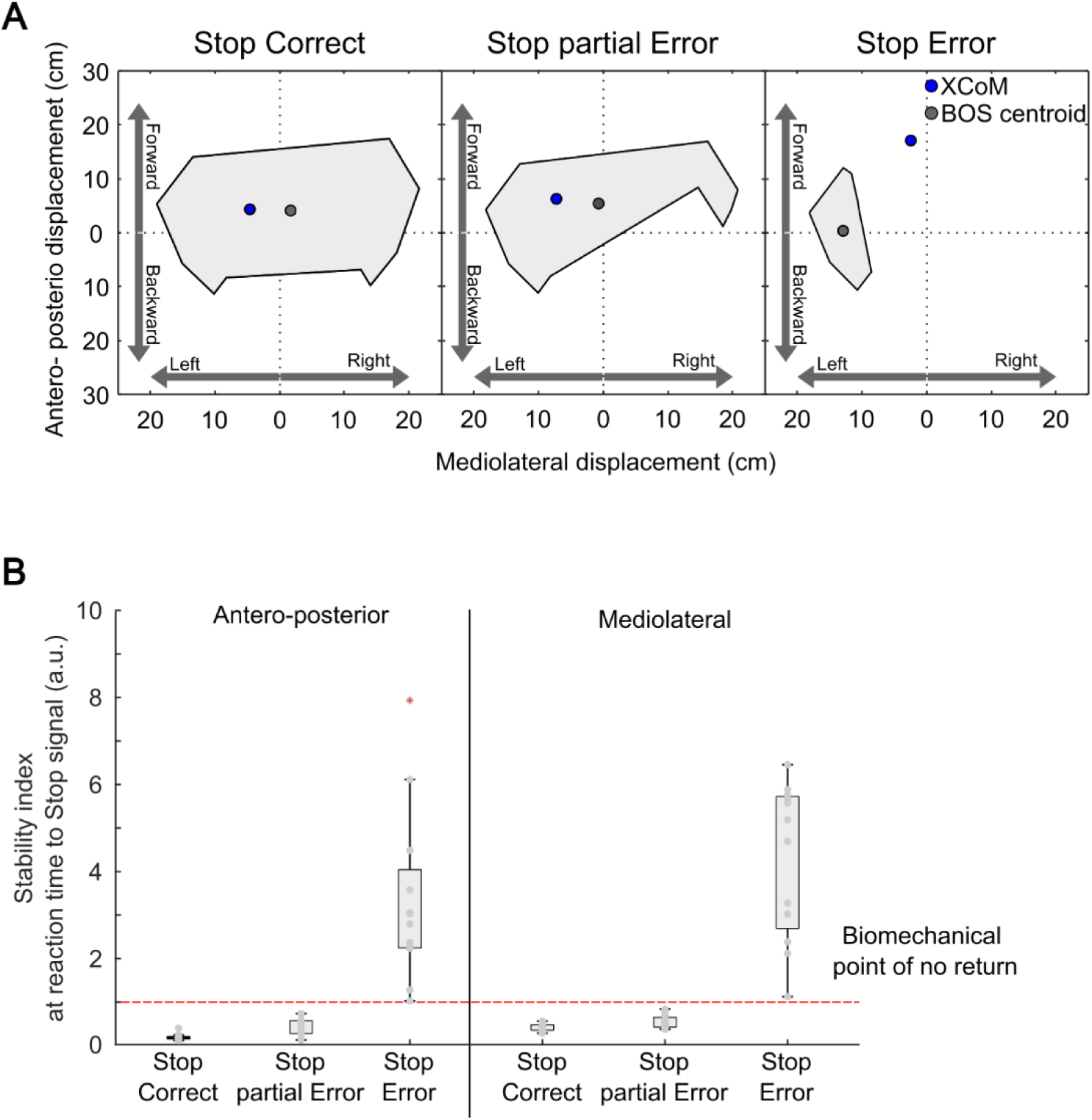
Relationship between the BOS and COM at the time of the reaction to the Stop signal. A. Antero- posterior and mediolateral displacement of COM at the time of stop reaction time. B. Stability index calculated at the Reaction time to Stop signal in the three different categories of Stop trials. One way Anovas (antero-posterior: **F**(1,2)=29.0, ***p***<0.001; mediolateral: **F**(1,2)=54.3, ***p***<0.001) and Bonferroni post-hoc comparisons (***ps***< 0.05) revealed a significant difference between the Stop error trials and both Stop correct and Stop partial error trials. Significant differences between the latest were not detected (***p***s>0.05).

## Discussion

In the present work, we implemented a gait initiation stop signal task to investigate how complex movement could be integrated in the framework of the stop signal paradigm. By extracting a set of variables from the monitoring of APA preceding the first step onset, we illustrate a scenario in which all detected variables— including the latency of APA reaction time to the Go and Stop signal, as well as the Stop Signal Delay duration—potentially influence step inhibition. Specifically, we observed that any combination of the detected variables that allows for the correction of APA will result in successful step inhibition. The only requirement is that the projection of the center of mass must be kept within the boundary of the base of support. Thus, we found that step inhibition is possible within a point of no return determined by the biomechanical state of the body.

The great majority of studies on motor control have used experimental settings where relatively simple movements as key press or reaching to a peripheral target ^37^. Similar experimental settings have been employed for studying motor inhibition ^16,19,22,31^. However, all ignored to evaluate the contribution of the postural adjustments. Indeed, several of our daily actions ^3,4,38^, require a simultaneous adjustment of the posture for allowing the proper completion of a movement. Recently APA have been described during reaching movements in crouched body posture ^5^. In the present study, we focused on step initiation, a complex behavior requiring a timely coordinated sequence of postural adjustments ^6,39^. While previous studies have addressed how the processing of external stimuli can affect the temporization of APA before starting a step ^10,11,13,14,40,41^, our focus here is on how APA are modified when an external event leads to an efficient cancellation of the prepared gait initiation. By referring to the theoretical assumptions underlying the stop signal paradigm, here we outlined a framework that well embedded the possible behavioral outcomes occurring when the cancellation of a prepared step is required. Firstly, here we demonstrated that theoretical assumptions of the horse race model, developed from the study of simpler movements, well fitted such a complex motor behavior. Within this framework, we found that trials with longer APA reaction times to the Go signal were more likely to be interrupted, particularly when a Stop signal was presented after a short SSD, and the APA Reaction time to the Stop signal was rapid. Crucially, we noticed that successful movement inhibition only occurred when these events aided in adjusting the movement until the COM did not cross a biomechanical threshold. Within this range of adjustment, all countermanding commands worked to maintain the body in a stable starting position. This was observed in Stop Correct and partial Error trials. In the former, the longer APA reaction time to the Go signal, as detected in the displacement of COP and CPM variations coupled with an earlier occurrence of the Stop signal at a shorter SSD, facilitated movement cancellation with minimal perturbation to body stability and kept the COM within the base of support boundaries. This correction was also aided by a short Reaction time to the Stop signal.

In Stop partial Errors, the COP and COM reacted earlier to the Go signal, and the Stop signal occurred at a longer SSD. However, the rapidity of the APA Reaction time to the Stop in these trials helped maintain the initial body position despite a perturbation that caused heel lift. As in previous trials, successful inhibition occurred because, at the time of the Reaction to the Stop signal, the COM’s projection fell within the edges of the BOS. Finally, in Stop Error trials, the earlier COP reaction time to the Go signal and the occurrence of the Stop signal at a longer SSD were not counteracted by the APA Reaction time to the Stop duration. In fact, in these cases, the longer APA Reaction time to the Stop duration caused a deviation to the COP trajectory, which was reflected in the COM trajectory when it moved well outside the BOS boundaries. In this scenario, the only way to prevent falls was to complete the initiated step.

Using a similar approach, Kwag et al. (2024) identified a relationship between the onset time of the center of mass (COM) and the timing of the stop signal, which influenced the probability of successfully inhibiting gait initiation. Their study, conducted on a large population of young and older participants, involved presenting a small number of stop trials (only 9 out of 36 total trials) with a fixed stop-signal delay (SSD).

Our findings complement and expand upon theirs by testing participants with a significantly larger number of trials (90 stop trials out of 300 total) presented randomly, with SSDs adjusted based on individual performance. This approach enabled us to obtain a measure of reaction time to the stop signal that aligns with the theoretical framework of the stop-signal paradigm. The limited number of trials used in Kwag et al. (2024) may have restricted their ability to fully assess the adherence of their results to the established knowledge on motor inhibition derived from the stop-signal paradigm ^17^.

The present results fit previous findings, emphasizing a sequential progression of events leading up to the observable initiation of a movement, such as a key press. This progression starts with neuronal activity programming the movement and culminates in the engagement of muscular activity ^20,21,27^. In Figure 4 we present a schematic that integrates our observation on APA with the theoretical background of the stop signal paradigm. This figure illustrates the dynamic interaction between the Go and Stop processes, which ultimately determine behavioral outcomes and contribute to estimating the Stop-Signal Reaction Time (SSRT) (as evidenced by the top reaction time distributions). Recent research (Jana et al., 2020; Ramawat et al., 2024; Raud et al., 2022) has shown that reaction times to both Go and Stop signals, based on movement onset, tend to overestimate the actual response to these signals, obscuring the temporal dynamics of the stop process and failing to provide insight into its completion time. Our results support the hypothesis that every Stop signal initiates the stop process, but its effectiveness in halting depends on the timing of its intervention and potential to rectifying the ongoing APA.

**Figure 4.**
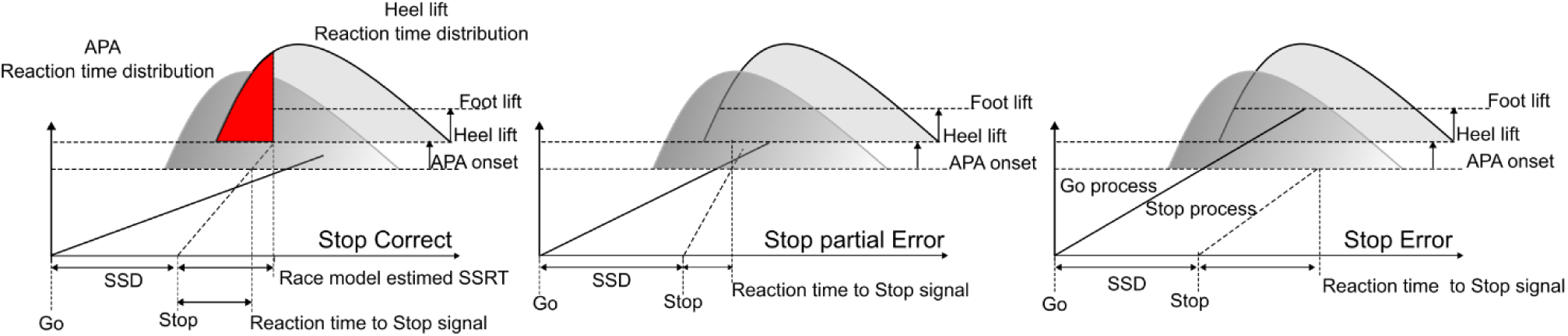
Biomechanical variables at play in step inhibition. Time evolution of the Go and Stop process in relation to the APA onset and Heel lift Reaction time distributions. In Stop correct trials (Left panel) the Stop process passes the threshold corresponding to the APA onset first. The APA reaction time to the Go signal could be still detected but there is no generation of movements. In Stop partial Error trials (Middle panel), the Go process is more rapid and, therefore, could pass both the APA and Hell thresholds. However, the Stop process is still sufficiently rapid to allow for a correction and the step is aborted. In the Stop Error trials (Right panel), the stop process is too slow and cannot compete with the Go process passing all the thresholds and generating the full step.

### Implication for extending the stop signal paradigm to study complex movements and the underlying neural correlates

The present approach has yielded two significant achievements in the investigation of action inhibition using the Stop Signal Task. Firstly, we successfully detected important variables at the single-trial level, such as reaction time to the Go signal in Stop trials and the reaction time to the Stop signal in Stop Error trials. These variables were not previously identified in most studies even if their detection is crucial for evaluating the ongoing of movement’s initiation and cancellation. Tentative SSRT detection at the single-trial level successfully estimated this variable in ballistic and semi-ballistic movements. This estimation was achieved by monitoring the activity of agonist or antagonist muscles involved in the task ^20,21,31^. In the case of finger movements, the SSRT measurement was obtained by identifying partial error trials. These trials represented a subset of Stop Correct trials where there was an initial increase in agonist muscle activity after the Go signal, as for starting a movement, followed by a decrease in activity to maintain the finger in the starting position as a response to the Stop signal. These findings indicated the existence of a point of no return, beyond which movement cancellation was unlikely, and provided an estimate of the SSRT based on the onset of muscle activity attenuation ^20,21,42^.

Even in the present work we detected a subset of Stop partial Error trials that provided a reliable estimate of the SSRT. Here Stop partial Error trials were a proportion of Stop Error trials where the heel left the starting position, but the participant was still in the condition to refrain from movement generation. In addition, thanks to the monitoring of APA time evolution, here we were able to detect changes in velocity of the COP and COM trajectories that reliably estimated this variable also in Stop Correct and Stop Error trials. These measures, accounted for the hypothesis that only the attempts to cancel a programmed movement occurring before a point of no return can end up with a success. Importantly our results strengthen the hypothesis of a point of no return in movement control facing against the original hypothesis that correctly inhibited response in the SST were those corresponding to a partial preparation that never reached an overt manifestation even in absence of a stop signal ^27^. This evidence strengthens the plausibility of the original formulation of the race model assuming the stop process to be triggered every time a stop signal is presented and challenges the hypothesis that stop failure is accounted by the occurrence of “trigger errors” ^43,44^.

The trigger error hypothesis aligns with the race model’s characterization as a winner-takes-all process, suggesting that some error trials occur when the stop signal fails to trigger the stop process. While this perspective may be plausible for studying ballistic or semi-ballistic movements such as saccades or finger key presses/releases ^18,19,45^, it may not fully apply to non-ballistic movements as arm extension for which tentative of on-fly movement stopping are documented ^31^.

This limitation arises from the difficulty in monitoring intermediate steps between the generation of motor commands by the nervous system and the downstream dynamics of movement execution. Consequently, the original formulation of the race model reliably accounted for motor inhibition only in contexts where movement initiation coincided with its overt manifestation, as preceding events were not observable. In addition this model well fitted the approaches implemented for studying the neuronal computations going on in cortical and subcortical structures subtending both ballistic ^18,28,30,46^ and non-ballistic movements ^23,47^.

According to this model, the reaction time to the stop signal (SSRT) can be obtained by the speed of the Go process. When the Go process is excessively fast, it can override the Stop process, resulting in a movement despite the stop command. By analyzing the fastest reaction times in the distribution—those quick enough to escape the stop signal—the model provides an estimate of the average SSRT across a block of trials (illustrated by the red area in Figure 4, leftmost panel). This SSRT estimate has been instrumental in investigating motor inhibition across various contexts and clinical conditions ^17^. However, relying solely on an average SSRT provides an incomplete picture of motor inhibition, potentially obscuring specific features of different types of stop trials. Our findings reveal distinct differences in reaction times to the stop signal among correct, partial error, and error trials. These patterns, observed in healthy individuals, could vary in patient populations and offer valuable insights into specific aspects of motor inhibition.

Notably, even in simple movements with semi-ballistic dynamics, such as finger movements, attempts at motor inhibition have been detected during movement execution, after muscle recruitment. Correctly stopped trials often exhibit partial electromyographic (EMG) responses, while stop error trials display weaker EMG activity compared to Go trials ^20,35,36^. Consistent with these findings, we observed partial gait initiation in partial error trials and shorter steps in stop error trials compared to Go trials (Figure 1D). Furthermore, we identified reaction times to the stop signal that further support the conclusion that the Stop process was not interrupted solely by the Go process reaching a threshold.

Another important parameter to consider is the threshold level, which can be arbitrarily set. In classical approaches, only the overt detection of movement is considered, allowing for the definition of a threshold preceding the effective start of the movement. This threshold could align with the recruitment of muscle activity or, as in our study, with the onset of anticipatory postural adjustments (APAs). For example, by setting the threshold for distinguishing between correct and error stop trials at the point when the stepping foot completely leaves the ground (see Figure 4), we could define additional thresholds marking earlier stages of movement initiation. These thresholds likely reflect different levels of motor plan processing by the nervous system, from central to peripheral structures ^45,48,49^.

The present results reflect what recently observed in the dynamic of the neuronal activity in the Dorsal premotor cortex of monkey performing a reaching version of the SST. In that case, a movement was not initiated until the dynamics of the neuronal network was maintained within the boundary of a given configuration. Movements started only when the neuronal trajectory crossed this boundary ^24^. Here, we observed at a behavioral level that the relation between the body kinematics and the BOS marked a boundary between a control and a ballistic state for this behavior. Accordingly, models of APA control hypothesized a role in of premotor areas in programming and rectifying this set of postural adjustments (see ^48,49^).

Our findings plausibly indicate that after having integrated biomechanical variables influencing the body stability in models of motor control it will be achieved a reliable understanding of environmental interaction in contexts closer to real life situations ^16,50^. Understanding the roles of these variables in action inhibition is crucial for investigating the role played by each node of the neural circuits subtending action inhibition as the frontal cortex and premotor cortex ^23,32,50^ or the cerebellum ^51–53^ . The importance of proprioceptive feedback influencing postural control ^8^, as well as the managing of these afference by the spinal cord ^48,49^, subcortical nervous structure ^54,55^ or the somatosensory cortex ^8,48,49^ will enrich the understanding of the neural circuit managing complex behavioral adjustments.

To conclude, we obtained direct measurements of critical variables involved in postural adjustments during the initiation of steps that fit within a consolidated theoretical model of motor control. Considering the opportunity to monitor online variations of these variables, we draw the possibility of predicting the outcome of complex movements. These results provide a solid framework for designing and testing tools for accident prevention and the rehabilitation of neurological patients with posture control impairments, such as those suffering of Parkinson’s disease (Yitayeh and Teshome, 2016).

## Limitations of study

Despite the high number of trials completed by the participants, the small sample size might limit the generalizability of our findings. While our within-subject design provides enough power to detect significant effects, the results may not be representative of the broader population. This means that extending our conclusions to a broader population requires further testing.

Additionally, the individual differences among the 12 participants could introduce variability that would be otherwise controlled in a larger sample. Focusing on a limited age range also prevents the extension of our results to a wider population. To address all these points, further tests are necessary.

## Acknowledgments

The authors are grateful to Dr. Isaac Kurtzer for valuable discussions.

## Author contributions

L.F. collected and analyzed the data. L.F., S.R, I.B.M., V.R, F.D. and A.R. discussed the results and commented on the manuscript. E.B., P.P., S.F., F.D. and A.R. contributed to the study design. E.B., L.F., S.F. wrote the manuscript.

## Use of AI technology

During the preparation of this work the authors used ChatGpt AI technology for grammatical purposes. After using this tool/service, the authors have reviewed and edited the content as necessary and take full responsibility for the content of the publication.

## METHODS

### Participants

Twelve healthy participants (4 women, 8 men) aged 33.3±7.1 years, with an average weight of 69.3 ±17.0 kg, and an average height of 169.7±8.2 cm, were recruited for the study. None had orthopedic or neurological issues or ongoing drug therapies. They provided informed consent. Participants completed a preliminary test using a revised Edinburgh Handedness Inventory (EHI) to determine preferred starting limb for walking (Oldfield, 1971). The study adhered to the Declaration of Helsinki and was approved by the local Ethics Committee (N. 0078009/2021). Data collection took place at the movement analysis laboratory, Department of Occupational and Environmental Medicine, Epidemiology and Hygiene, INAIL, research center in Monte Porzio Catone, Rome.

### Gait initiation stop signal task

Each participant was initially positioned on two out of eight dynamometric platforms, with their feet parallel and bare, aligned with the movement analysis lab’s x-axis following the International Society of Biomechanics’ standard ^56^. Subsequently, a contact sensor was inserted between the platform and the heel of the participant’s preferred limb, while a computer monitor was positioned 2 meters away and 0.7 meters high for visual stimulus presentation. Participants received instructions to maintain focus on the stimulus monitor and initiate forward movement as quickly as possible. Before the experiment, participants underwent practice trials to familiarize themselves with the setup. Each gait initiation trial commenced when the participant’s heel made contact with the sensor, triggering the stimulus system to display a traffic light signal on the monitor’s center. This signal was then replaced, after a variable delay (randomly between 1 and 1.5 seconds), by a forward-facing arrow (595x822 pixels) acting as a Go signal.

In 70 % of trials required participants to initiate forward movement (Go trials). The initiation was determined by the release of sensor-heel contact, marking the participant’s Heel lift time (RT). Go trials with an RT exceeding one second or occurring before the Go stimulus were considered ineligible.

In 30% of trials (Stop trials), the forward-facing arrow was replaced by a stop road sign (1024x1024 pixels), serving as a Stop signal. Upon presentation of the Stop signal, participants were instructed to halt their motor response and remain stationary. The Stop signal followed a variable delay from the Go signal (Stop Signal Delay; SSD), determined using a scaling algorithm based on participants’ performance. The SSD, initially set at 50 ms, was adjusted automatically, increasing by 50 ms after each successfully cancelled Stop trial and decreasing by 50 ms following an uncanceled Stop trial, as per the algorithm. A trial was deemed a Stop Error trial if the participant lifted their heel from the contact sensor after the Stop signal; otherwise, it was a Stop Correct trial. Participants received auditory feedback on the correctness of their response. Each participant completed 10 blocks of 30 trials (300 trials in total), with SSD values maintained across consecutive blocks. Before the data collection each participant completed a block of 30 practice trials.

## INSTRUMENTATION

In this experimental study, two systems were combined into one to capture both the behavioral and motor patterns linked to gait initiation. One system focused on movement analysis, collecting kinematics, dynamics, and muscle activity data, while the other managed behavioral aspects, handling visual and auditory stimuli and recording participant responses. These systems were synchronized to create an integrated and timed dataset. Only kinetic and kinematic data were analyzed in this study.

### Behavior instrumentation

The SST paradigm was controlled by a PC running Windows XP, a monitor (18”), and a loudspeaker system. An algorithm implemented in the MATLAB computing environment managed the projection of visual stimuli on the screen and acoustic stimuli on the speakers, as well as the acquisition and transmission of information about both the emitted stimuli and the behavior of the participants via the sensor under the heel via a parallel port.

### Gait Initiation recording instrumentation

A stereo-photogrammetric motion analysis system with optoelectronic technology was used to collect the kinematics data (SMART-DX 6000 system: BTS, Italy, Milan). Eight infrared cameras with a sampling rate of 340 Hz were employed, as well as 32 reflective markers were placed above the anatomical reference points ^57,58^. In detail, the markers were placed above on the cutaneous projections of spinous processes of the seventh cervical vertebra, tenth dorsal vertebra, vertebra sacral and bilaterally on the frontal and parietal bones, inion, acromion, lateral humeral condyles, radial processes, ulnas styloid, third metacarpal, superior anterior iliac spine, and above the lower body bilaterally on the cutaneous projections of large trochanter, lateral femoral condyle, fibula head, lateral malleoli, head of the third metatarsus and heels (Figure S1 A).

This whole-body marker placement protocol, as well as the large number of markers used, were chosen to obtain a measurement of whole-body kinematics. This decision was made to be able to use the segmental method to calculate the Center of Mass (COM; method explained later), which is evidently one of the most accurate methods for its estimation(Gutierrez-Farewik et al., 2006; Ranavolo et al., 2017) .

The kinetics data of the ground reaction forces (GRFs), with their center of pressure, were acquired using eight dynamometer platforms (Kistler 9286B; Kistler, Winterthur, Switzerland) at a sample rate of 680Hz (Figure S1 B). Each platform was made up of four load cells, with capacitive technology, each of which was placed in one of the platform’s four corners. The peculiarity of these eight platforms was that they were virtually interconnected, so that the system could be considered as a single force platform during data extraction.

## DATA ANALYSIS

3D reconstruction software (SMART Tracker and SMART Analyzer: BTS, Italy) and MATLAB (R2019b 9.7; MathWorks, USA) were used to process the kinematics and kinetics.

### Classification of Stop partial Error and Error trials

During data collection, we identified two types of Stop Error trials. In one type, participants lifted both the heel and the toe, completing the movement. These were classified as stop error trials. In the other type, only the heel was lifted without advancing the limb forward. Similar to the finger versions of the SST, these trials initially began as errors but were corrected during the trial, resembling partial-stop error trials^20,21^, which were analyzed separately from regular stop trials.

To biomechanically characterize Stop partial Errors, we applied a 5 cm threshold to the forward displacement of the malleolus marker:

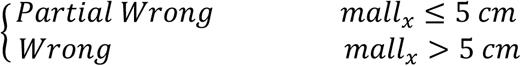

This threshold was determined in a data-driven manner, based exclusively on the observation of the collected data.

### Kinematic and kinetic pre-processing

To obtain an ordered and homogeneous collection of all Gait Initiation data, only a temporal window from 0.2 s before the Go signal until 1.6 s after it was studied. This time window ensures a thorough examination of APAs in all the studied participants’ behavioral responses.

To keep only the signals of interest, all raw kinematics and kinematics data were first low-pass filtered with a zero-lag fifth-order Butterworth filter with a 10 Hz frequency of cut off.

### Computation of the center of mass and the center of pressure

The pre-elaborated kinematics data were then used to evaluate the total body COM by combining them together with the subjects’ anthropometric data and the estimate of body segment parameters^61,62^.

The computation was carried out by treating the COM as the centroid of a set of *n* body segments and computed as follows:

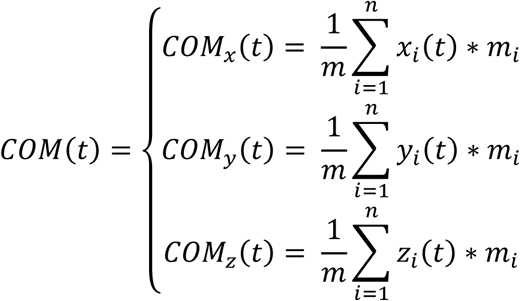

Where, *COM*_*x*_(*t*), *COM*_*y*_(*t*) and *COM*_*z*_(*t*) are the instantaneous components of the *COM*(*t*) position along the *x* (anteroposterior direction), *y* (mediolateral direction) and *z* (vertical direction) axes respectively; *m* is the mass of the system under consideration (weight of the subject), n is the total number of body segments under consideration (*n* = 12; head, trunk, 2 arms, 2 forearms, 2 hands, 2 legs, and 2 feet); while *x*_*i*_(*t*), *y*_*i*_(*t*) and *z*_*i*_(*t*) are the coordinate in the space of the *i-th* segment, and mi is its mass.

The COP position was calculated from the distribution of the GRFs on the platforms as the point location of the ground reaction force vector and was directly released by motion analysis system, and computed as follows:

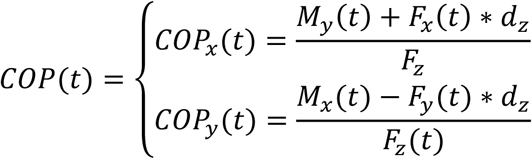

where *COP*_*x*_ (*t*) and *COP*_*y*_ (*t*) represent the instantaneous location of the *COP*(*t*) on the force platform along the axes *x* (anteroposterior direction) and *y* (mediolateral direction), respectively; *M*_*x*_(*t*) and *M*_*y*_(*t*) are the moment of the force platform along *x*-axis and *y*-axis; *F*_*x*_ (*t*), *F*_*y*_ (*t*) and *F*_*z*_(*t*) represent the force in *x*-axis, *y*-axis, and *z*-axis, respectively, and *d*_*z*_ represents the height of the top of plate above measurement plane (*xy*).

### Trial-by-trial detection of reaction time to the Go and Stop signal from the time evolution of COM and COP trajectories

A template-based algorithm was developed to automatically recognize the reaction times to Go and Stop stimuli from COM and COP trajectories. This approach is widely utilized in the literature to detect motor patterns in motion analysis signals, including motion capture, inertial sensors, and surface electromyography^63–66^ .

In this study, the template model was computed and applied to the velocity profiles of the COM and COP. These velocities were obtained by calculating finite difference derivatives and applying a low-pass filter. Specifically, a zero-lag third-order Butterworth filter with a cut-off frequency of 5 Hz was used to smooth the signals.

### Template-based detection of reaction time to Go and Stop signal

To assess each participant’s reaction to the Go signal, marking the onset of anticipatory postural adjustments (APAs), two templates were created: one for the COP trajectory and another for the COM trajectory, regardless of trial type. These templates were generated by averaging five randomly selected trials from the Go trial dataset, all aligned to the Heel-off event. For each of the five trials, the velocity was calculated, and the resulting signals were averaged. Each averaged signal was then appropriately trimmed to account baseline variations at the signal’s onset. This template was subsequently used to determine the reaction time to the Go signal across all Go and Stop trials.

For Stop trials, three distinct templates were created for each trajectories (COM and COP) to capture the reaction time to the Stop signal: one for Stop Correct, one for Stop partial Error, and one for Stop Error trials. Each template was generated by averaging three randomly selected trials from each respective trial type, aligned to the Stop signal. The resulting averages were trimmed to account for signal baseline variations.

Supplemental Figure 2 illustrates the templates generated for detecting reaction times to Go and Stop signals in the COM trajectories for the three types of Stop trials. The algorithm applied to COP trajectories follows the same procedure but is not shown.

Reaction times to the Go and Stop signals for APAs were identified as the time points corresponding to the minimum Euclidean distance between the previously described templates and the recorded signals (Supplemental Figure 2B), calculated as follows:

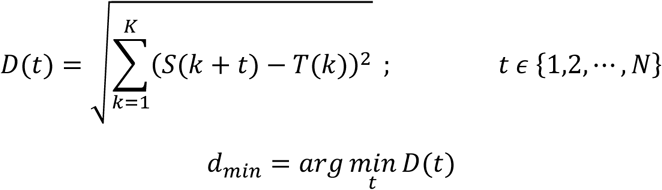

Where *S* and *T* are the signal and template respectively, *K* is the length of template and *N* is the length of signals.

### Stability index

Gait stability is a fundamental aspect of movement analysis, commonly studied by examining the relationship between the projection of the center of mass onto the ground (XCoM) and the base of support (BoS). Stability is achieved when the XCoM lies within the BoS, and it increases as the XCoM approaches the center of the BoS. These indices are frequently employed in gait analysis to evaluate stability during double support phases ^67,68^ or at gait termination ^69^. However, they are not typically applied to transient phases of limb progression, as investigated in this study.

To address this limitation, we developed a specific stability index, the Stability Index (STi), tailored for this experimental study. This index was computed based on the two fundamental parameters, XCoM and BoS, to assess stability during the phases of movement analyzed. STi is calculated as the distance between the BOS centroid and the XCoM along two directions: Anterior-posterior (x-axis) and Mediolateral (y-axis):

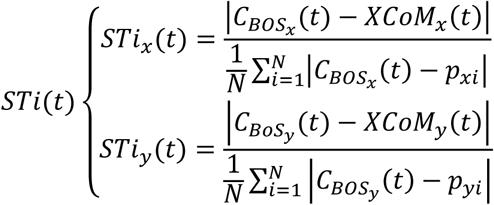

where *XCoM*_*x*_(*t*), *XCoM*_*y*_(*t*), *C*_*BOSz*_(*t*) and *C*_*BOSx*_(*t*) represent the instantaneous anteroposterior and mediolateral XCoM and centroids of the BOS positions, respectively.*p*_*xi*_ and *p*_*yi*_ are the vertices defining the perimeter of the BOS, and *N* is the number of vertices composing the BoS perimeter (*N* =11).

The Stability Index (STi) quantifies the relative position of the XCoM with respect to the BoS:

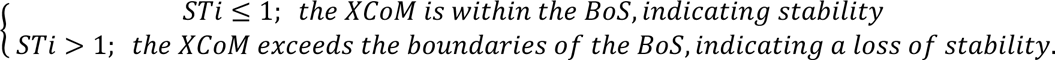

The XCoM is a concept derived from a linearized inverted pendulum model, which can be considered as a point on the floor at a distance from the COM directly proportional to the COM velocity. It was calculated using the following formula:

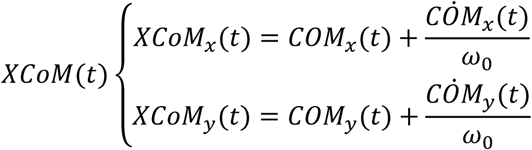

where *XCoM*_*x*_(*t*), *XCoM*_*y*_(*t*) and *COM*_*x*_(*t*), *COM*_*y*_(*t*) represent the instantaneous anteroposterior and mediolateral XCoM and COM positions, respectively;, *COM*_*x*_(*t*) and *COM*_*y*_(*t*) are the instantaneous anteroposterior and mediolateral CoM velocities; and ω_0_ is the natural frequency, calculated using the following equation:

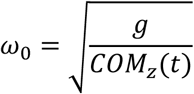

where *g* is the acceleration of gravity and *COM*_*z*_ is the effective height of the body’s COM above the floor.

The BoS, which generally refers to the area beneath a person’s feet and includes every point on the supporting surface, was computed in this study using foot markers and anthropometric measurements collected in the initial phase of each subject’s trial (Supplementary Figure 3A). More specifically, the perimeter of the BoS area was defined instant by instant by the outermost foot markers (Supplementary Figure 3B). Each marker was included or excluded from the set of available markers for BoS calculation based on its elevation relative to the baseline: markers with an elevation smaller than 5 mm were included, while those greater than 5 mm were excluded.

### Foot distance during swing phase

To evaluate the distance traveled by the foot during the swing phase at the gait initiation, we calculated the displacement of the foot’s center of mass, previously determined (see Center of Mass section). Specifically, the distance was computed as the difference between the initial and final positions of the center of mass trajectory during the swing phase.

### Estimate of the Stop Signal Reaction Time according to the horse race model’ assumptions

The mathematical approach used in this research project to estimate the SSRT was the integration method^15^. In depth, this method estimates the point at which the stop process finishes by ‘integrating’ the RT distribution and finding the point at which the integral equals probability of Stop Error trials. The finishing time of the stop process corresponds to the nth RT, with *n* = *number of RTs* in the RT distribution of Go trials multiplied by *p*(*Stop Error*).

## STATISTICAL ANALYSIS

A repeated-measures ANOVA was performed to examine significant differences in APA reaction times to the Go and Stop signals, measured from the COM and COP, across SSD and STi (anteroposterior and mediolateral directions), with trial type as a within-subjects factor. For the APA reaction times to the Go signals, four conditions were analyzed: Go, Stop Correct, Stop Partial Error, and Stop Error. For the APA reaction times to the Stop signals, SSD, and STi, three conditions were compared: Stop Correct, Stop Partial Error, and Stop Error. A Bonferroni post-hoc analysis was conducted to pinpoint specific differences between trial types.

Additionally, a paired two-sample t-test was used to evaluate the difference in step distance between Go trials and Stop Error trials.

A post-hoc statistical power analysis using G*Power (3.1.9.7) ^70^ was performed for ANOVA repeated measures, within factors, with α =0.05, based on partial eta squared (η^2^_*p*_) calculated by F ^71^. The analysis indicated that the power (1-β err prob) for each ANOVA was > 0.95. The significance level was set to 0.05.

More specifically, we assessed the adequacy of the sample size (12 participants) for SSRT and RT analyses. For SSRT (1 group, 3 measurements, correlation = 0.76), simulating an effect size of 0.4 with α = 0.05 yielded a power of 0.98. For RT (1 group, 4 measurements, correlation = 0.85), the same parameters yielded a power of 0.99. An additional a priori power analysis for ANOVA (1 group, 4 measurements, assumed correlation = 0.70) confirmed a power of 0.95 for detecting an effect size of 0.4 with α = 0.05.

A correlation analysis (Pearson coefficient) was performed between each APA reaction time to the stop signal measured from COM, COP, and the SSRT estimated with the integration method.

## Supplemental Information

**Figure S1.**
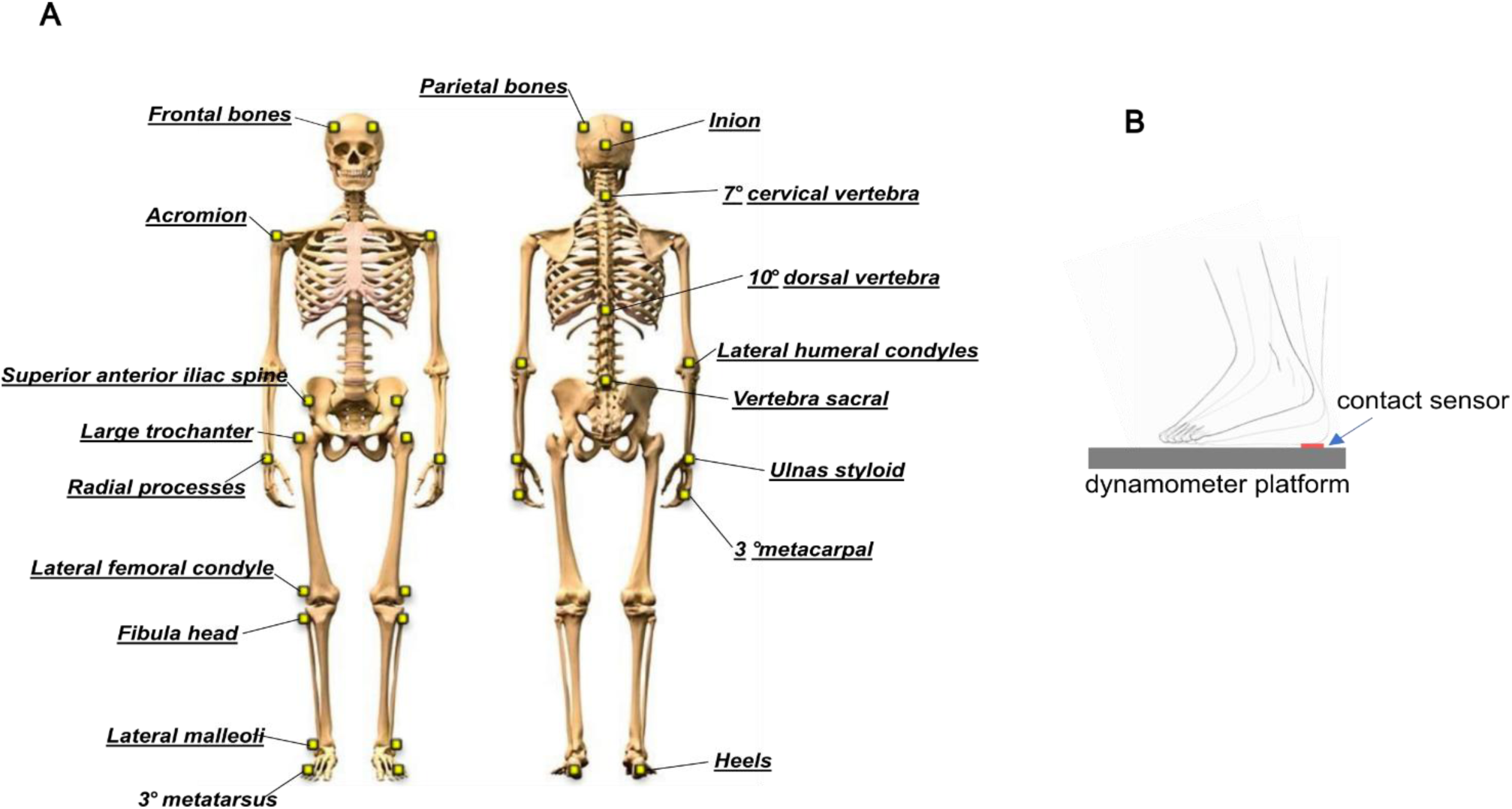
Schematic of devices for detection of behavioral, kinetic, and kinematic variables. A, displacement of kinematic sensors on the body target points. B, Relationship between foot, contact sensor and force detection platform.

**Figure S2.**
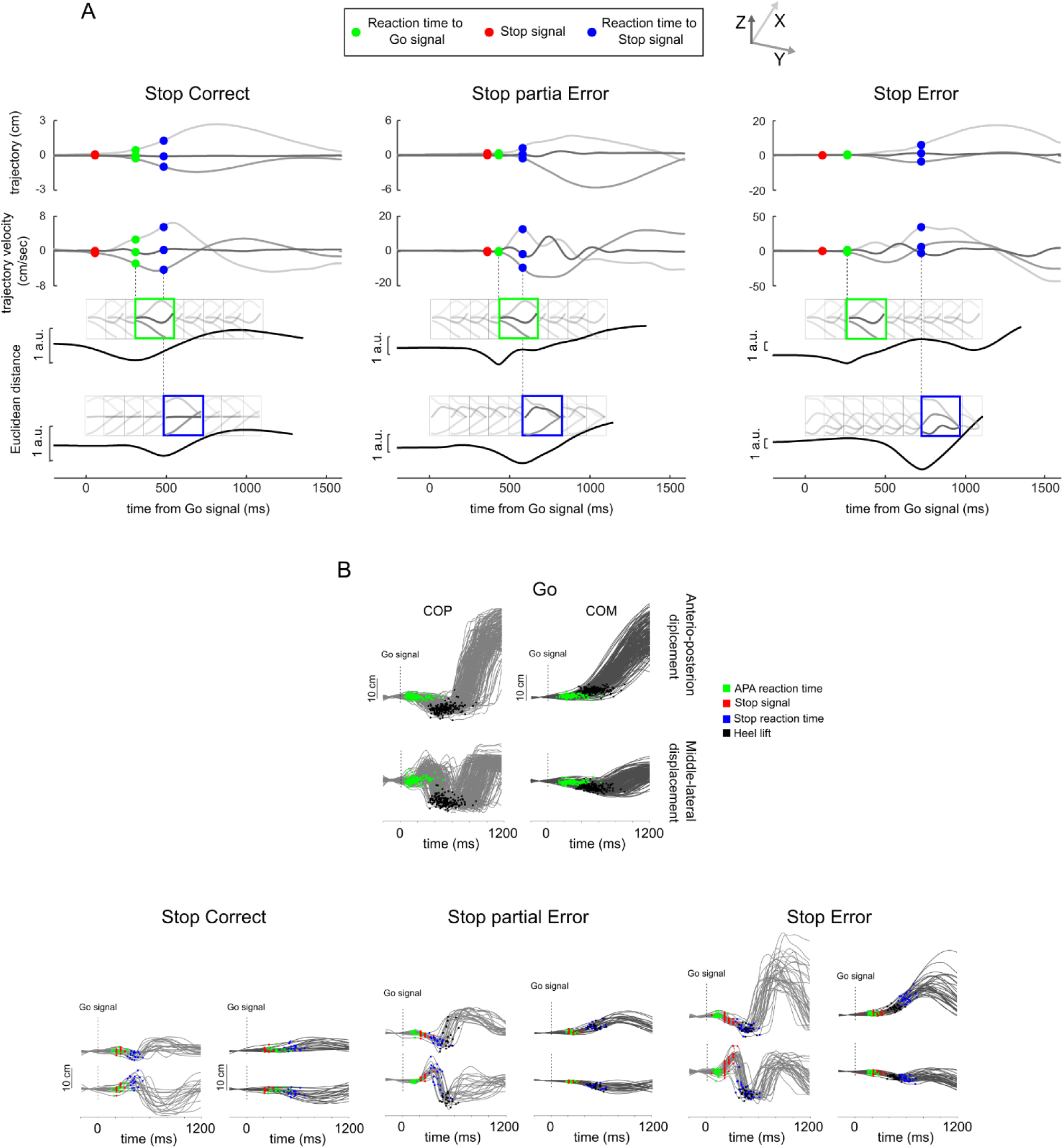
Single trial-based computation of reaction time to Go and Stop signal in Stop trials on COM. A Upper row over time displacement of the X, Y and Z component of COM. Bottom rows velocity of X, Y and Z components of COM for each type of stop trial. The figure highlights the Euclidean distance between template of the pattern of change in velocity when participants reacted to the Go (Green rectangles) and Stop (Blue rectangles) signal. The reaction time to the Go and Stop signal is taken as the minimum Euclidean distance between the template and the velocity tracks (green and blue dots). B) time course of the Antero-posterior (upper plots) and (Mediolateral) shifting of a single trial of the four possible outcomes in the task the COP (left column) and COM (right column) of a participant performing the task. Each line corresponds to a single trial of the four possible task’s outcomes. Green squares and circles mark for each trial the time of APA onset obtained by the COP and COM respectively by applying the method illustrated in A.

**Figure S3.**
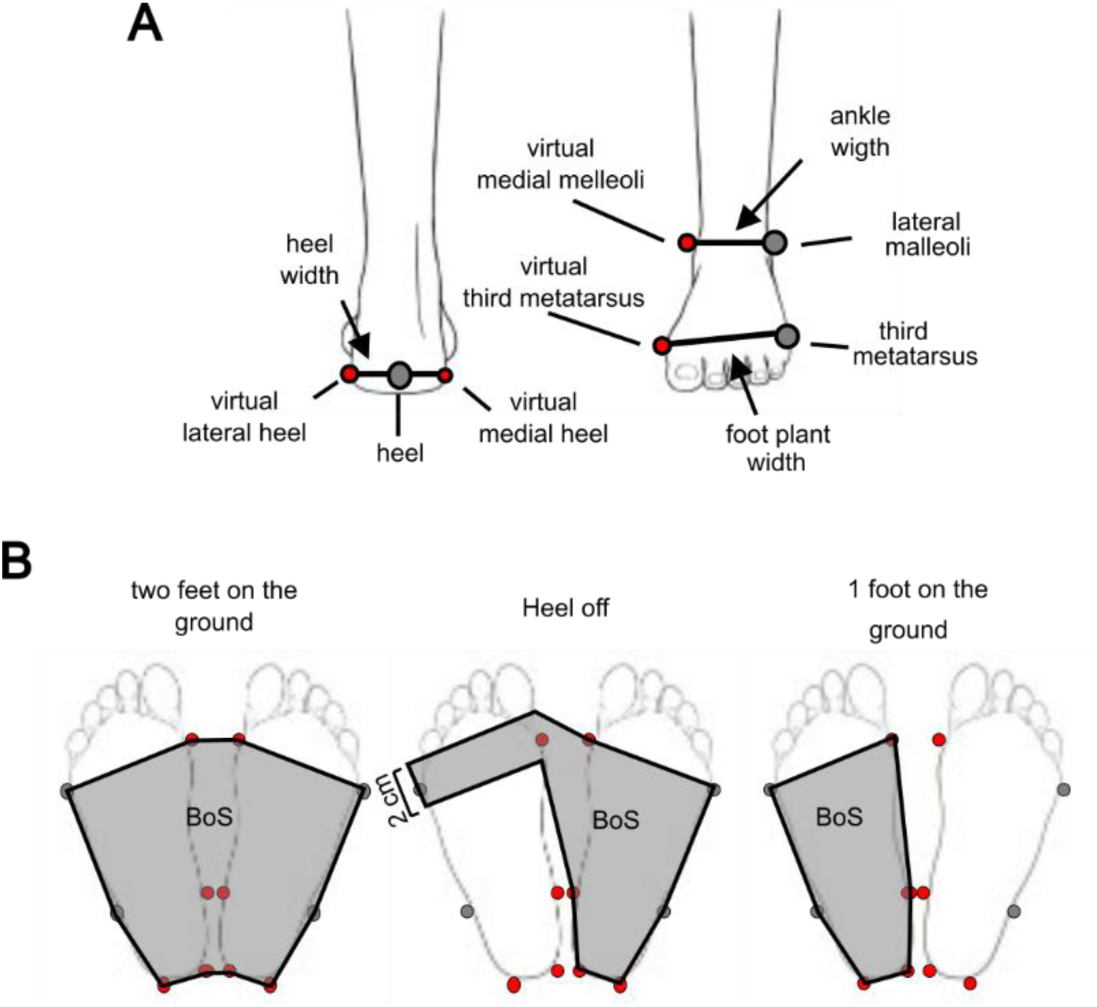
Computation of base of support. A, Anthropometric measures used for the computation of subjects’ base of support. B, Three possible scenarios during gait initiation and their supporting bases are depicted schematically: 1) BoS when participants have both feet on the ground (leftmost panel); 2) BoS when participants are in the heel-off phase (middle panel); 3) BoS when participants have only one foot on the ground (rightmost panel).

